# Network supporting contextual fear learning after dorsal hippocampal damage has increased dependence on retrosplenial cortex

**DOI:** 10.1101/209866

**Authors:** Cesar A.O. Coelho, Tatiana L. Ferreira, Juliana C.K. Soares, João R. Sato, Maria Gabriela M. Oliveira

**Affiliations:** Departamento de Psicobiologia, Universidade Federal de São Paulo – UNIFESP, São Paulo, SP, Brazil, zipcode 04023064; Centro de Matemática, Computação e Cognição, Universidade Federal do ABC, UFABC, São Bernardo do Campo, SP, Brazil, zipcode 09606070

## Abstract

Hippocampal damage results in profound retrograde, but no anterograde amnesia in contextual fear conditioning (CFC). Although the content learned in the latter have been discussed, the compensating regions were seldom proposed and never empirically addressed. Here, we employed network analysis of pCREB expression quantified from brain slices of rats with dorsal hippocampal lesion (dHPC) after undergoing CFC session. Using inter-regional correlations of pCREB-positive nuclei between brain regions, we modelled functional networks using different thresholds. The dHPC network showed *small-world* topology, equivalent to SHAM (control) network. However, diverging hubs were identified in each network. In a direct comparison, hubs in both networks showed consistently higher centrality values compared to the other network. Further, the distribution of correlation coefficients was different between the groups, with most significantly stronger correlation coefficients belonging to the SHAM network. These results suggest that dHPC network engaged in CFC learning is partially different, and engage alternative hubs. We next tested if pre-training lesions of dHPC and one of the new dHPC network hubs (perirhinal, Per; or disgranular retrosplenial, RSC, cortices) would impair CFC. Only dHPC-RSC, but not dHPC-Per, impaired CFC. Interestingly, only RSC showed a consistently higher centrality in the dHPC network, suggesting that the increased centrality reflects an increased functional dependence on RSC. Our results provide evidence that, without hippocampus, the RSC, an anatomically central region in the medial temporal lobe memory system might support CFC learning and memory.

**AUTHOR SUMMARY:** When determined cognitive performances are not affected by brain lesions of regions generally involved in that performance, the interpretation is that the remaining regions can compensate the damaged one. In contextual fear conditioning, a memory model largely used in laboratory rodents, hippocampal lesions produce amnesia for events occurred before, but not after the lesion, although the hippocampus is known to be important for new learning. Addressing compensation in animal models has always been challenging as it requires large-scale brain mapping. Here, we quantified 30 brain regions and used mathematical tools to model how a brain network can compensate hippocampal loss and learn contextual fear. We described that the damaged network preserved general interactivity characteristics, although different brain regions were identified as highly important for the network (e.g. highly connected). Further, we empirically validated our network model by performing double lesions of the hippocampus and the alternative hubs observed in the network models. We verified that double lesion of the hippocampus and retrosplenial cortex, one of the hubs, impaired contextual fear learning. We provide evidence that without hippocampus, the remaining network relies on alternative important regions from the memory system to coordinate contextual fear learning.

## INTRODUCTION

Lesion studies examine primarily the extent to which the brain can compensate the damaged region. For instance, a post-lesion impaired behavioral performance means that the remaining collective brain regions do not compensate the damaged region (Geschwind, 1965; Aggleton, 2008). In contextual fear conditioning (CFC), hippocampal lesions result in profound retrograde amnesia (of pre-lesion events), but no anterograde amnesia (post-lesion events; Frankland et al., 1998; Wiltgen et al., 2006), suggesting that new learning is supported by the reminiscent regions. Evidence for hippocampal participation in CFC acquisition have been provided by manipulations ranging from pharmacological injections such as muscarinic (Gale et al., 2001) and NMDA receptors blockade (Schenberg and Oliveira, 2008), to optogenetic approaches (Liu et al., 2012). Thus, although hippocampus participates in CFC learning if it is functional during acquisition, CFC learning can occur after hippocampal loss.

The hippocampal loss compensation in CFC inspired cognitively-oriented hypotheses about the content learned by non-hippocampal regions, some proposing a fragmented (elemental) context representation (Nadel and Willner, 1980; Rudy, 2009), others proposing a still configural representation (Fanselow, 2010). These hypotheses further propose that hippocampus has preference over the non-hippocampal regions. This accounts for the impaired CFC observed after temporary manipulations, during which hippocampus inhibits the non-hippocampal regions while unable form long-term memory (Fanselow, 2010). However, little attention has been given to the compensatory regions. Although parahippocampal cortices were pointed out as putative candidates (Rudy, 2009), the regions involved in hippocampal loss compensation in CFC have not been empirically addressed. Investigating how these regions learn and store CFC information can help to understand the dynamics of hippocampal function and its interactions within the memory systems.

There is evidence for a large number of regions to compose the neural circuits involved in CFC (Fanselow and Poulos, 2005; Maren, 2011) and spatial/contextual memory (Bucci and Robinson, 2014). Understanding compensation of a lesion requires assessing complex interactions among the remaining regions and their possible changes. Network approaches assess complex brain interactions based on the representation of elements (i.e. brain regions, neurons) and connection concepts (i.e. projections, functional connectivity), and offer quantitative tools for a data-driven assessment of network characteristics related to brain structure and function (Bullmore and Sporns, 2009).

Large-scale network studies based on structural and functional MRI data have been paving a solid ground in cognitive neuroscience (Medaglia et al., 2015; Mišić and Sporns, 2016). They have explored functional network topology in the brain (Achard et al., 2006) and its importance to learning (Bassett et al., 2011) and emotion (Kinnison et al., 2012). Network studies have also been useful in identifying crucial brain regions (hubs) for network function (van den Heuvel and Sporns, 2013), and to identify functional network changes after traumatic brain injuries (Hillary and Grafman, 2017) and in psychiatric disorders (Crossley et al., 2016; Sato et al., 2016). Some studies took advantage of rodent models and employed network analysis in the expression of the activity-dependent gene *c-fos* after remote CFC retrieval (Wheeler et al., 2013) and later empirically interrogated the network hubs given by the model (Vetere et al., 2017). Here, we used a similar rationale to investigate hippocampal compensation in CFC. We used the phosphorylated cAMP response element binding (pCREB, active form of CREB), which is critical to learning-induced synaptic plasticity (Alberini, 2009), as our marker of brain region engagement; and examined activation and coactivation of brain regions of hippocampectomized rats after a CFC session. Using network analysis, we examined how the compensatory network might support CFC learning and memory. We hypothesized that different network attributes in the ‘damaged network’ could be underlying hippocampal compensation in CFC learning. Further, we performed double lesions to empirically validate some results that indicated possible models for compensation of hippocampal loss in CFC (**Figure 1**).

**Figure 1:**
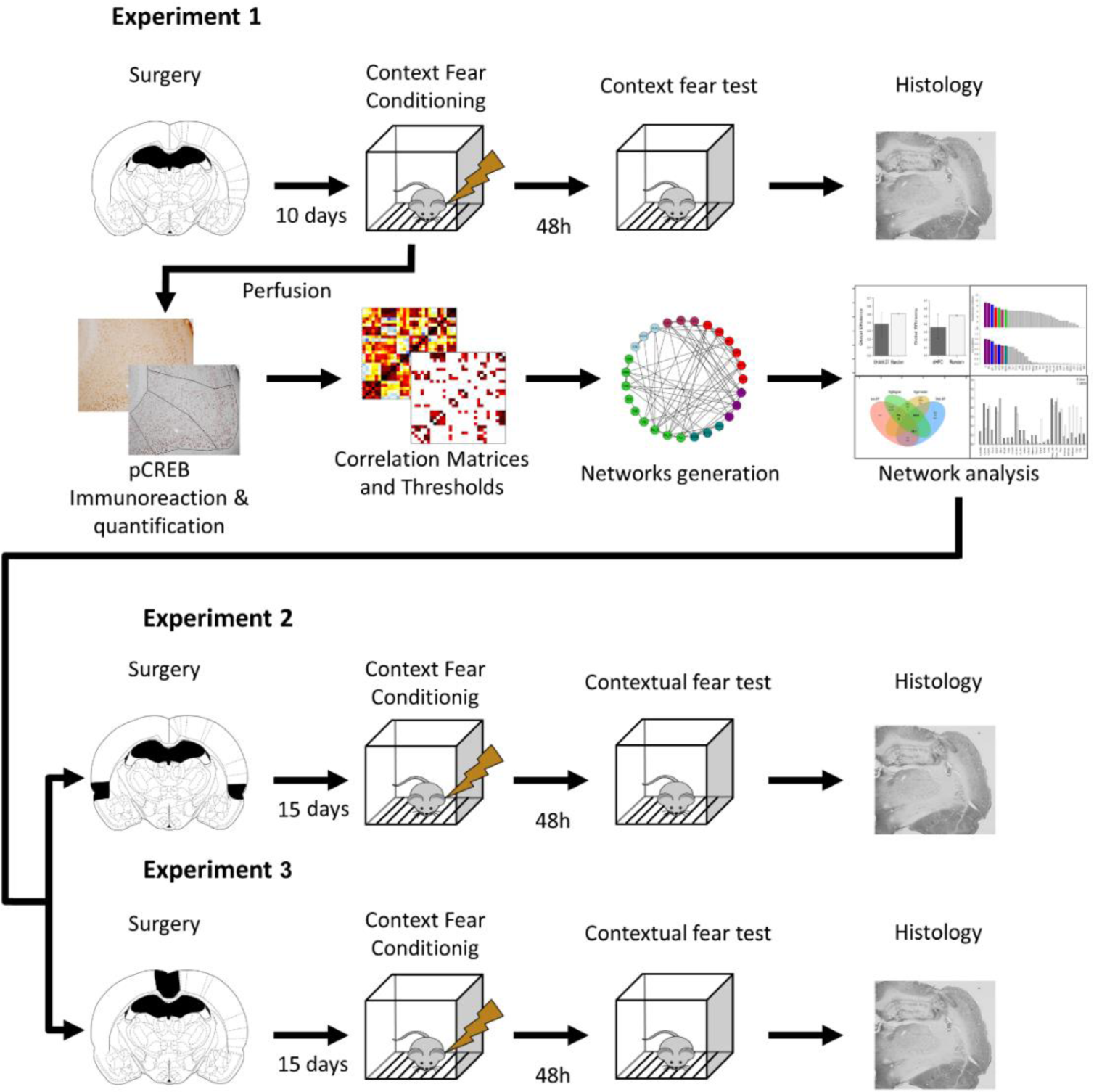
Overview of the experimental design. *In Experiment 1*, the rats underwent pre-training dHPC lesions and, after recovery, a contextual fear conditioning session. Half the sample underwent fear memory test 48 h later and half was perfused 3 hours after CFC and had their brains processed and stained for pCREB protein. Thirty regions had their pCREB expression quantified and their pCREB inter-regional correlations computed. After thresholding the correlations, we analyzed the networks properties and compared them between the groups. Following network analysis, *Experiments 2 and 3* employed double lesions to test if the network differences observed could be empirically supported.

## RESULTS

### Experiment 1 – Network underlying contextual fear learning in the absence of dHPC

*Experiment 1* aimed to explore how CFC learning under dHPC damage changes other brain regions activity and interactivity patterns compared to CFC learning in normal rats. We compared pCREB expression levels between the groups and modelled functional networks based on pCREB expression correlations. Then we employed network tools to explore differences between damaged and control groups. In *Experiment 1*, the rats initially underwent bilateral electrolytic lesions in the dHPC or SHAM surgery. After surgical recovery, the rats underwent a CFC training session. Half the cohort was perfused 3 h after the training session, and their brains processed for pCREB immunolabelling. The other half was returned to the homecage and tested for contextual fear memory 48 h later. A group of immediate shock controls (Imm) was added to the cohort that was tested for contextual fear memory.

#### dHPC damage does not alter CFC memory

The histological examination of dHPC lesions revealed that the cellular loss was overall confined to the dorsal part of the hippocampus, with occasional lesion to the overlaying cortex due to the electrode insertion (**Figure 3a**). The cohort tested for contextual memory had the freezing behavior measured as memory index, and was compared among the groups. The sample size in the memory test cohort was 32 (SHAM: N = 12; dHPC: N = 12; Imm: N = 8). A bootstrapped one-way ANOVA showed a significant group effect (F_2,29_ = 8.822, p = 0.0011). Multiple comparisons performed with p-corrected t-tests showed higher freezing time in both SHAM (p = 0.0001) and dHPC (p = 0.0178) groups compared to Imm group, but not statistically different from one another (p = 0.4044; **Figure 3b**). A KS test confirmed these results, showing no difference between SHAM and dHPC samples (D_24_ = 0.3333, p = 0.2212) and both different from Imm group sample (SHAM: D20 = 0.917, p = 0.0001; dHPC: D_20_ = 0.667, p = 0.0070; **Figure 3c**). A Cohen’s d showed a medium effect size between SHAM and dHPC means (d = 0.630) and large effect sizes between these two groups and Imm (SHAM: d = 2.226; dHPC: d = 1.275). These results show no effect of dHPC lesion in CFC learning and are in agreement with past studies (Wiltgen et al., 2006).

**Figure 2:**
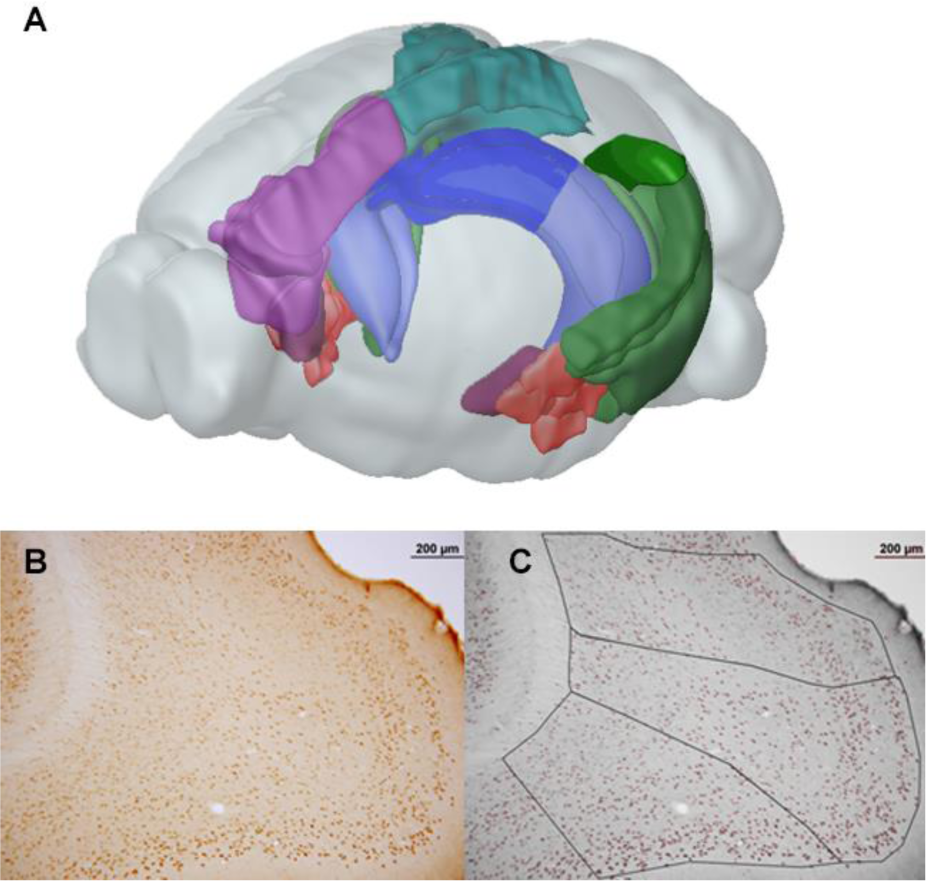
3D diagram of a rat brain showing the regions quantified for pCREB (A). Representative photomicrograph of a pCREB immunolabelled brian slice before (B) and after (C) nuclei quantification and region parcellation by Cellprofiler. The scalebars indicate 200 µm.

**Figure 3:**
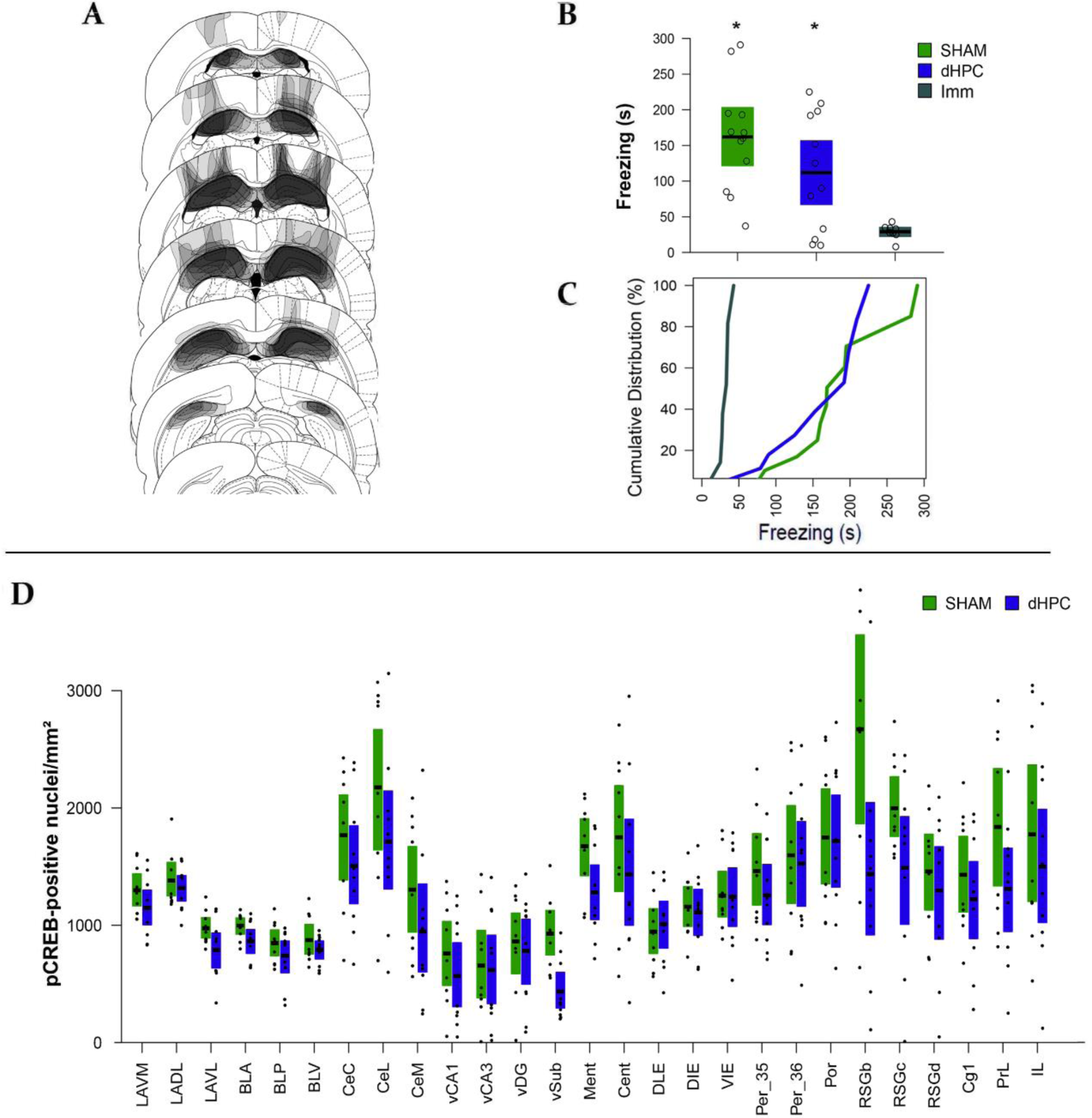
dHPC lesion does not impair CFC. (A) Schematic diagram showing the distribution of the lesions in the dHPC group. (B) Mean (black line) and bootstrapped 95% CI of the Total Freezing Time during the five min context fear memory test of dHPC (N = 12), SHAM (N = 12) and Imm (N = 8) groups. The open circles show data distribution in each group. (C) Cumulative distribution of the sample as a function of Freezing Time showing the sample distributions. The “*” shows a significant difference from Imm at level of p<0.05. (D) Mean (black line) and Bootstrapped 95% CI of the mean (boxplots) of the pCREB-positive nuclei density in each region and each group. The black dots show the data point distributions.

#### dHPC damage does not alter the overall pCREB levels in the quantified regions

In the pCREB immunolabelling cohort, we tested whether the dHPC lesion altered pCREB expression after CFC learning in any of the studied regions by comparing the pCREB expression in each region between dHPC and SHAM groups. The **Figure 3d** shows the pCREB expression in each region and each group. The sample size in the pCREB expression cohort was 19 (SHAM: N = 9; dHPC: N = 10). A visual inspection of the pCREB data reveals an expression roughly similar to previous studies (Stanciu et al., 2001; Trifilieff et al., 2006). We analyzed the pCREB-positive nuclei density by comparing each region between the groups using t-tests with bootstrap resampling. There was only one marginally significant difference showing a higher level in the SHAM group in the vSub (t = 3.699, fdr-corrected p = 0.053). All other regions did not present a significant difference. This result indicates that dHPC damage diminish the pCREB expression in the vSUB, but otherwise does not alter the overall pCREB-positive nuclei density compared to the SHAM group.

#### Functional Networks

We used the pCREB data to generate correlation-based networks for the SHAM and dHPC groups. As the SHAM groups has three regions absent in the dHPC group (dCA1, dCA3 and dDG), a third network was generated as “SHAM with no dorsal hippocampus”, SHAM-nH, to allow for direct comparisons between the networks (**Figure 4**). For each matrix, three networks were generated considering correlations with p-values under the threshold of 0.05, 0.025 or 0.01, respectively. The networks had a very similar connectivity density that, as expected, linearly decreased as the thresholds increased in rigor (SHAM networks had 92, 64 and 40 edges respectively, SHAM-nH had 74, 53 and 35 edges, and dHPC had 77, 53 and 32 edges). In all thresholds, the networks had one big connected component and 3-5 disconnected regions in the most stringent threshold (0.01). Although in our study negative correlations were included as absolute values in the edge weights, no negative correlations survived the thresholds. Overall, the networks presented some visual differences in their pattern of connectivity, which we formally tested in the analyses that follow.

**Figure 4:**
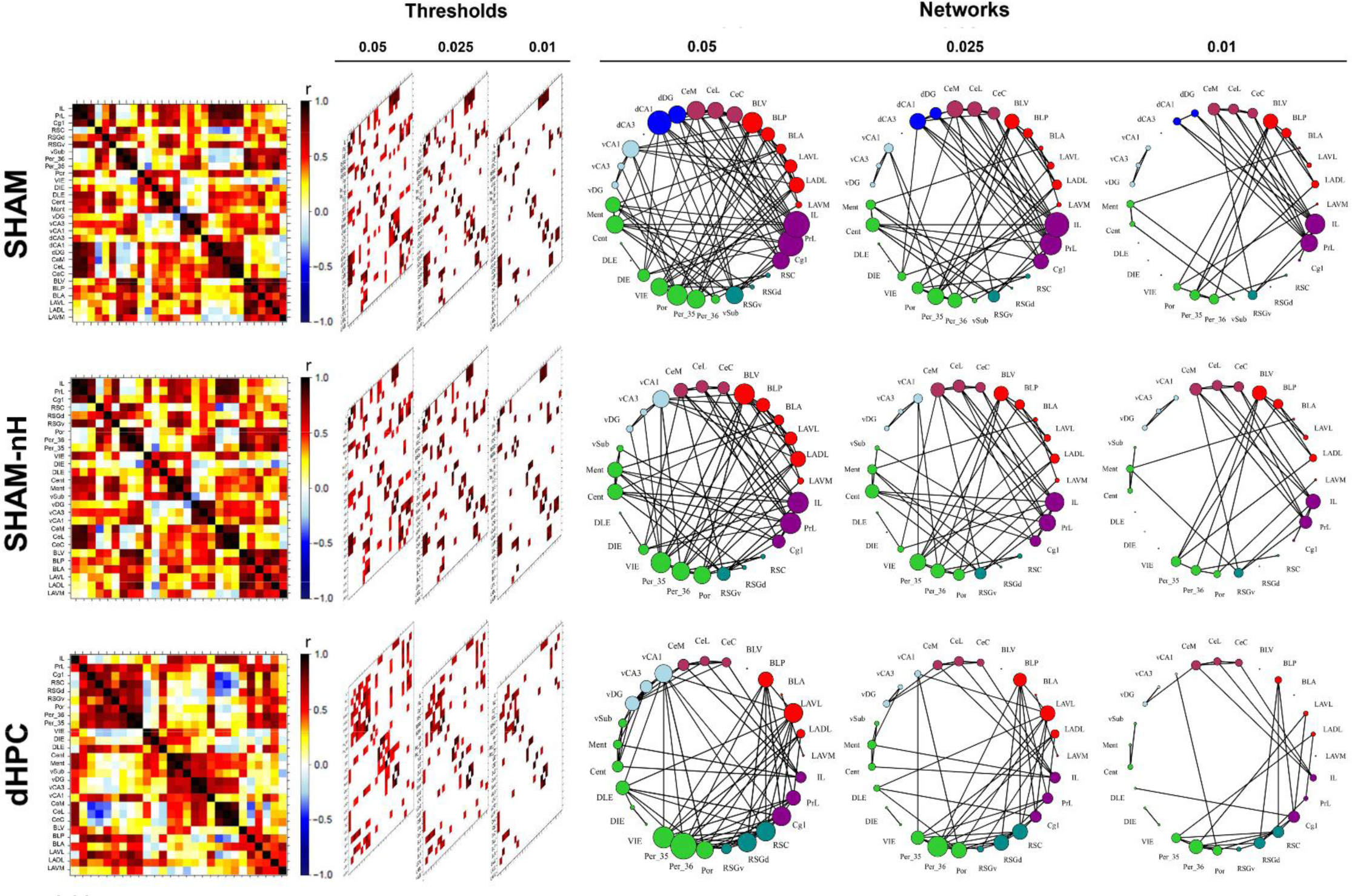
Generation of the connectivity networks in each group. After computing the inter-regional correlations (left), three thresholds were applied (p < 0.05, 0.025 and 0.01) and the most robust correlation coefficients (center) composed the networks (right). Networks were generated for SHAM (top), SHAM-nH (middle) and dHPC (bottom) matrices. In the matrices, colors reflect correlation strength (scale, right). In the network, the colors of the nodes are coded according to the Table 1, and the sizes of the nodes represent their degree (number of connections).

**Table 1:**
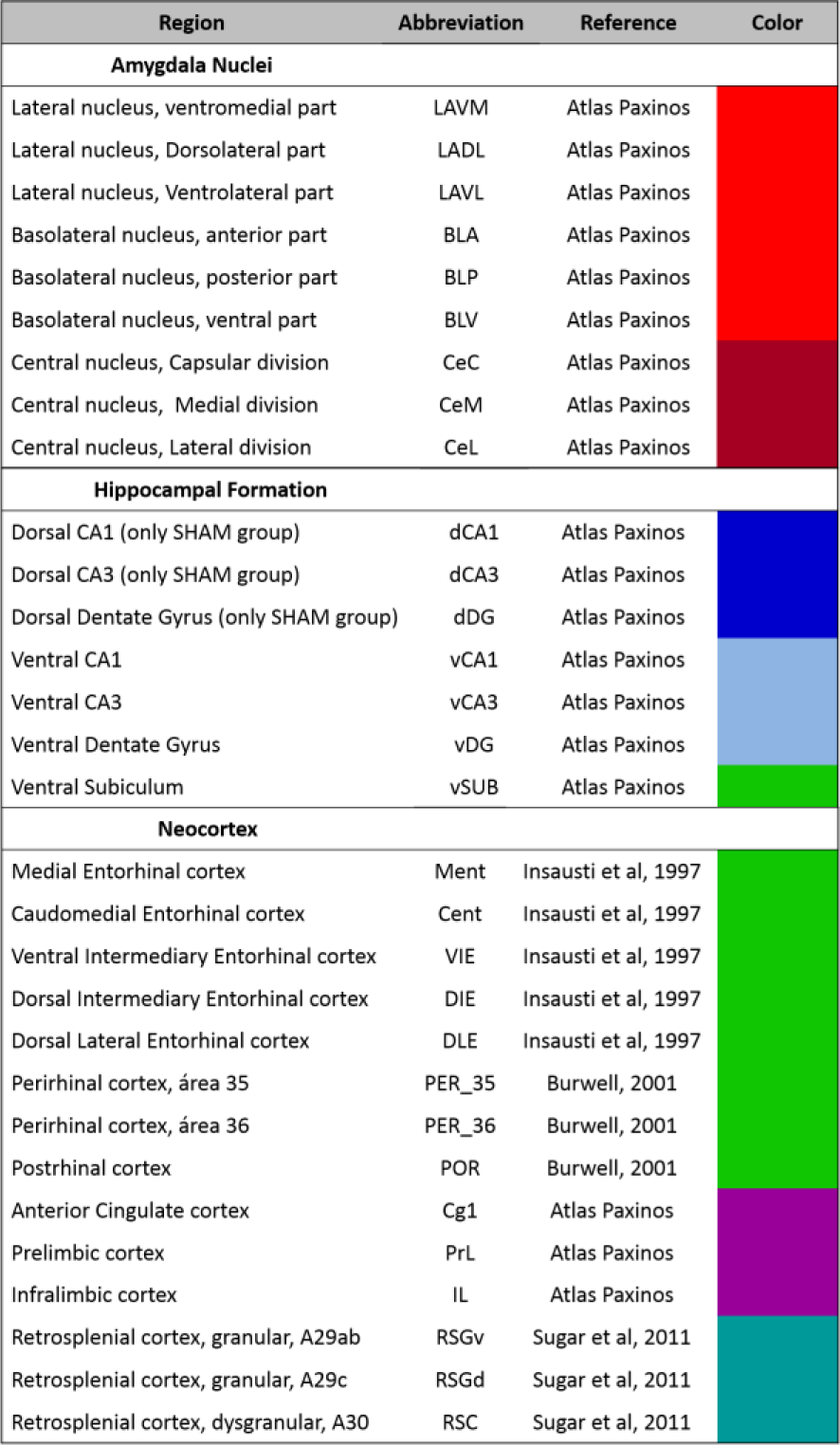
List of regions included in Experiment 1. The columns show the name of each region, the abbreviations and the source of the anatomical definition adopted, and the color code used in figures for each group of regions. Color code – Red: basolateral complex of the amygdala; Dark Red: Central Amygdala nuclei; Blue: dorsal Hippocampus; Light Blue: ventral Hippocampus; Green: parahippocampal regions; Purple: Prefrontal cortices; Magenta: Retroesplenial cortices.

#### dHPC damage did not alter small-worldness of the CFC learning network

We first tested whether the empirical networks (SHAM, SHAM-nH and dHPC) were small-world by comparing their global (Geff) and local (Leff) efficiencies to those of randomized null hypothesis networks. The **Figure 5** depicts the distribution of the empirical/randomized ratios of Geff and mean Leff for all networks and thresholds. In all cases, Geff ratios are roughly around 1, with a slight decay on the 0.01 threshold. Similarly, the mean Leff ratios are consistently above 1, with the mean and upper range of ratios increasing and the threshold increases in rigor. Equivalent integration (Geff) and robustly higher segregation (Leff) values in empirical networks compared to randomized networks is consistent with small-world networks accounts (Watts and Strogatz, 1998; Latora and Marchiori, 2001). These results suggest that the networks engaged in CFC learning are small-world, which is in agreement with a previous work showing small-world organization in CFC retrieval networks (Wheeler et al., 2013). Further, dHPC lesion did not seem to change the dHPC network small-worldness or its levels of Geff and mean Leff compared to the other networks, suggesting that the overall characteristic interactivity in the network was not affected.

**Figure 5:**
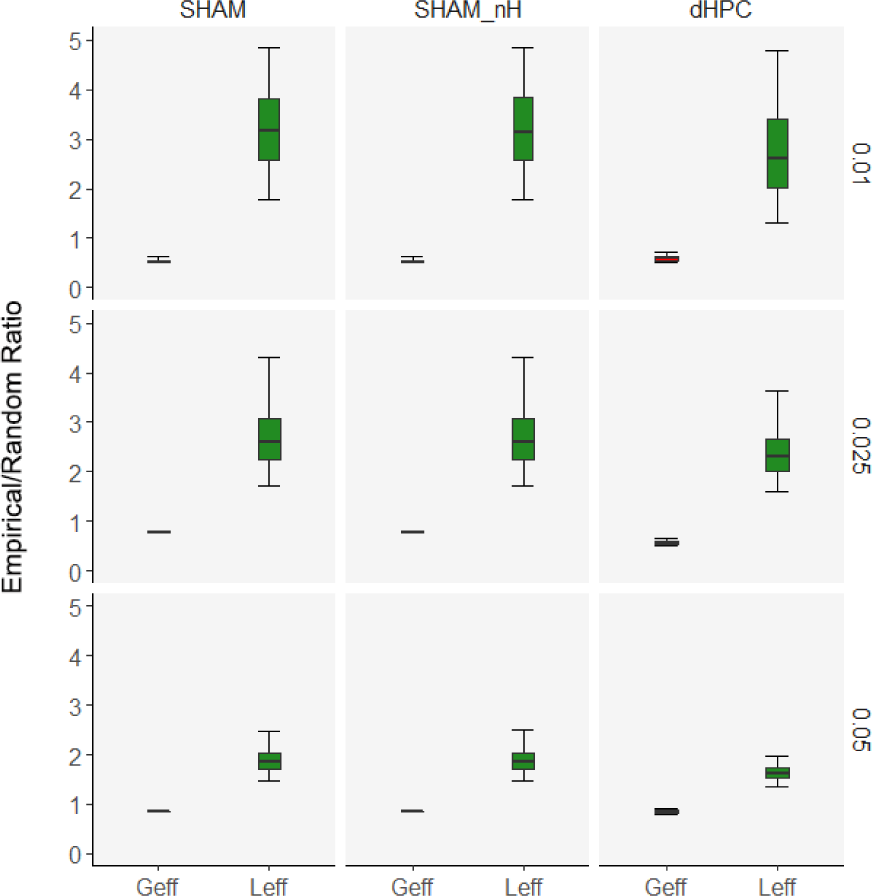
dHPC damage does not alter the CFC learning network small-worldness. Boxplots showing mean, lower and upper quartiles, and 95% CIs of the Empirical/Random Ratio of Geff and Leff for the SHAM (left), SHAM-nH (center) and dHPC (right) networks and on the 0.05(bottom), 0.025 (center) and 0.01(top) thresholds. Small-world networks are expected to have Geff ratios around 1 (empirical and randomized networks have roughly the same values) and higher Leff ratios (higher empirical values than those of the randomized networks).

#### dHPC network has alternative hubs

Hubs are defined as nodes positioned to confer the largest contributions to global network function, and are usually identified using multiple centrality metrics (van den Heuvel and Sporns, 2013). We considered as hub any region among the 25% most central regions in at least three of the four centrality metrics used (weighted degree, Wdg; eigenvector, Evc; closeness, Clo; and betweenness, Bet). Regions that were hubs across all thresholds were considered stable hubs. The **Figure 6a-b** shows the ranked centralities of each network and the metric intersections in each network for the threshold 0.05. In this threshold, the SHAM network showed the regions IL and BLV as hub, whereas in the SHAM-nH the BLV and Por were hubs, and the dHPC network the hubs were the Per_36, Per_35, RSC and LAVL. The **Figure 6c** shows which regions were considered stable hubs across the thresholds, in each network. In the SHAM network, the IL was the only region stably identified as a hub across all thresholds. In the dHPC network, the RSC, and the Per_36 were stable across all thresholds, and in the SHAM-nH network, no hub was stable across the three thresholds, but the IL was the closest region (hub in the 0.025 and 0.01 thresholds), similar to the SHAM network. Employing connection-based and distance-based metrics to identify a hub makes more likely that the identified well-connected regions are also inter-region or inter-modular connectors. Noticeably, the dCA1 was in the upper quartile of both connection-based metrics, but not the distance-based ones, across the all thresholds (not shown). These results suggest that different hubs emerged in the dHPC network. However, as the identification was descriptive, with no hypothesis test, it does not allow *a priori* interpretations regarding differences in the hub score between the networks. However, they are a first indication that there might be differences in the connectivity patterns between the SHAM and dHPC networks, as different regions emerged as hubs in these networks.

**Figure 6:**
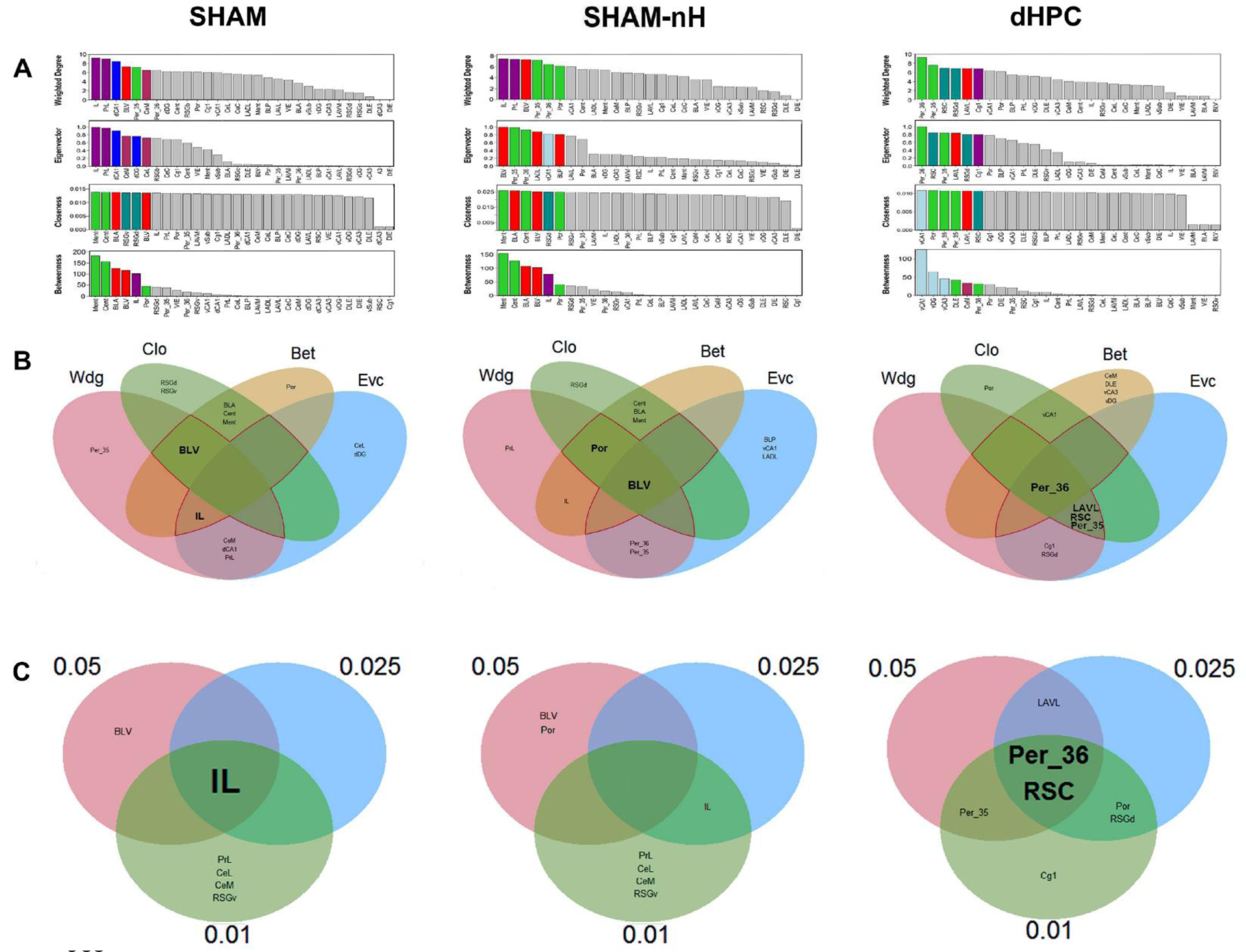
Hub identification of the networks. (A) The rankings of each centrality are shown for the (left) SHAM, (center) SHAM-nH and (right) dHPC networks under the 0.05 threshold. The colored nodes are the upper 25% most central in each metric. (B) The intersections of the upper 25% most central regions of each metric are shown for each network under the 0.05 threshold. Any region within the overlapping area of at least three metrics was considered a network hub (inside the red perimeter). The hubs were identified in the 0.025 and 0.01 threshold networks as well, and the hubs of each threshold were intersected (C) to verify the stability of the hub identification throughout the thresholds. Nodes are colored according to the code in Table **1**.

#### dHPC network hubs are associated to increased centrality measures

We addressed the hub score differences more formally and quantitatively by directly comparing the centralities between the groups in each region and each threshold using permutation test. The Table **2** resumes the results of the permutation tests for each region, metric and threshold. Most importantly, we observed that the identified stable hubs were overall associated with significantly higher centrality levels in some metrics, comparing the dHPC SHAM-nH networks. In the dHPC network, the RSC showed significantly higher Wdg and Evc in all thresholds, and the Per_36 showed higher Evc levels in the 0.025 and 0.01 thresholds, compared to SHAM-nH network. In the SHAM-nH network, the IL showed higher Evc levels in the 0.025 and 0.01 thresholds, compared to the dHPC network. Besides the stable hubs, some of the single-threshold or two-threshold hubs were also associated to significantly different centrality levels between the networks. In the dHPC network, the RSGd presented a higher Evc across all thresholds and a higher Wdg in the 0.025 and 0.01 thresholds. The LAVL had a higher Evc in the 0.025 threshold. In the SHAM-nH network, the BLV presented a higher Wdg across all thresholds, higher Bet in the 0.05 and the 0.01 thresholds, and higher Evc in the 0.05 threshold. Further, the CeM and PrL showed higher Evc, and the RSGv showed higher Bet, all in the 0.01 threshold. Some significant differences were present in non-hub regions such as BLP, vCA1, DLE and Por (higher metrics in dHPC network), and LAVM, BLA, and Por (higher metrics in the SHAM-nH network; Table 2). Lastly, some single-threshold hubs did not show significantly different centrality metrics in the thresholds they were considered hubs, such as LAVL, Per_35, Por and Cg1 (dHPC network) and CeL, Por (SHAM-nH network). These results provide evidence that when comparing SHAM-nH and dHPC networks, stable hubs in one network were associated to higher centrality levels relative to the other, and vice-versa. These data suggest that the CFC learning network under dHPC lesion has an increased dependence on its new hubs.

**Table 2:** Centrality Comparison between SHAM-nH and dHPC networks. The comparison is done for each region, centrality metric and threshold. Values in each cell show the permutation test p-value for each comparison. Green dots show significantly higher values in SHAM-nH, and blue dots show significantly higher values in dHPC (p < 0.05). In each threshold, green lines indicate SHAM-nH network hubs for that threshold, and blue lines indicate dHPC network hubs. Values with lines and dots in the same color show hubs associated with significant difference. Wdg: Weighted Degree; Evc: Eigenvector; Clo: Closeness; Bet: Betweenness.

|  | 0.05 |  |  |  | 0.025 |  |  |  | 0.01 |  |  |  |
| --- | --- | --- | --- | --- | --- | --- | --- | --- | --- | --- | --- | --- |
|  | Wdg | Evc | Clo | Bet | Wdg | Evc | Clo | Bet | Wdg | Evc | Clo | Bet |
| LAVM | 0.2943 | 0.1077 | 0.6353 | 0.7733 | 0.2893 | 0.5256 | 0.3666 | 0.0039 | 0.2019 | 0.2508 | 0.1355 | 1.0000 |
| LADL | 0.2210 | 0.0304 | 0.7423 | 0.8304 | 0.7016 | 0.0802 | 0.4052 | 0.7360 | 0.2547 | 0.1545 | 0.1666 | 0.1226 |
| LAVL | 0.3715 | 0.6279 | 0.7660 | 0.9015 | 0.1228 | 0.0155 | 0.4176 | 0.7357 | 0.8251 | 0.0831 | 0.1722 | 1.0000 |
| BLA | 0.1983 | 0.0844 | 0.6558 | 0.0314 | 0.3969 | 0.3737 | 0.3764 | 0.0146 | 0.2398 | 0.3772 | 0.1456 | 1.0000 |
| BLP | 0.8330 | 0.4279 | 0.7503 | 0.9500 | 0.4758 | 0.0214 | 0.4161 | 0.8981 | 0.3963 | 0.0728 | 0.1804 | 0.4444 |
| BLV | 0.0247 | 0.0001 | 0.6617 | 0.0180 | 0.0371 | 0.6883 | 0.3896 | 0.1367 | 0.0081 | 0.3326 | 0.1593 | 0.0482 |
| CeC | 0.5504 | 0.3965 | 0.7631 | 0.6283 | 0.4236 | 0.0791 | 0.4034 | 0.5262 | 0.2718 | 0.0896 | 0.1898 | 1.0000 |
| CeL | 0.6243 | 0.3717 | 0.7659 | 0.9603 | 0.5065 | 0.0761 | 0.4039 | 0.9338 | 0.3714 | 0.1107 | 0.1901 | 1.0000 |
| CeM | 0.6169 | 0.3369 | 0.7592 | 0.2273 | 0.2737 | 0.0608 | 0.3974 | 0.4616 | 0.3696 | 0.0419 | 0.1891 | 0.1555 |
| vCA1 | 0.8538 | 0.2397 | 0.7835 | 0.0238 | 0.4767 | 0.0839 | 0.4140 | 0.2549 | 0.3764 | 0.1778 | 0.1281 | 1.0000 |
| vCA3 | 0.3879 | 0.1898 | 0.8089 | 0.1097 | 0.8728 | 0.6306 | 0.4204 | 0.6615 | 0.3107 | 0.4872 | 0.2066 | 1.0000 |
| vDG | 0.2080 | 0.1553 | 0.8052 | 0.1018 | 0.4958 | 0.6294 | 0.4197 | 0.5935 | 0.9430 | 0.4923 | 0.2058 | 0.1753 |
| vSub | 0.6232 | 0.5306 | 0.7466 | 0.6393 | 0.9001 | 0.1529 | 0.3923 | 0.4696 | 0.6635 | 0.5091 | 0.1591 | 1.0000 |
| Ment | 0.3793 | 0.3053 | 0.7472 | 0.0028 | 0.7291 | 0.1700 | 0.3848 | 0.0078 | 0.2355 | 0.2487 | 0.1623 | 0.0752 |
| Cent | 0.4660 | 0.2796 | 0.7394 | 0.0086 | 0.2232 | 0.0487 | 0.3836 | 0.0118 | 0.9624 | 0.4101 | 0.1597 | 0.1925 |
| DLE | 0.0387 | 0.0304 | 0.8523 | 0.1882 | 0.2196 | 0.1065 | 0.4234 | 0.4952 | 0.2382 | 0.1103 | 0.0566 | 1.0000 |
| DIE | 0.1495 | 0.4349 | 0.8349 | 0.3036 | 0.1865 | 0.5832 | 0.9398 | 0.4403 | 0.0961 | 0.4487 | 0.1195 | 1.0000 |
| VIE | 0.1654 | 0.2881 | 0.8074 | 0.3417 | 0.2527 | 0.1057 | 0.4093 | 0.3023 | 0.2648 | 0.5224 | 0.1593 | 1.0000 |
| Per_35 | 0.8360 | 0.3149 | 0.7539 | 0.7491 | 0.5330 | 0.0816 | 0.4107 | 0.8274 | 0.7152 | 0.0870 | 0.1796 | 0.2965 |
| Per_36 | 0.1635 | 0.4930 | 0.7516 | 0.6756 | 0.1786 | 0.0200 | 0.3980 | 0.1187 | 0.1850 | 0.0453 | 0.1771 | 0.3251 |
| Por | 0.9951 | 0.5611 | 0.7315 | 0.5671 | 0.3084 | 0.0213 | 0.3751 | 0.0036 | 0.3161 | 0.0554 | 0.1693 | 0.1963 |
| RSGv | 0.5753 | 0.2276 | 0.7451 | 0.4060 | 0.3403 | 0.1777 | 0.3923 | 0.3584 | 0.2301 | 0.2210 | 0.2157 | 0.0213 |
| RSGd | 0.0570 | 0.0056 | 0.7404 | 0.3056 | 0.0434 | 0.0134 | 0.4072 | 0.8319 | 0.0240 | 0.0294 | 0.1178 | 0.0587 |
| RSC | 0.0123 | 0.0236 | 0.7635 | 0.5285 | 0.0304 | 0.0103 | 0.3886 | 0.6506 | 0.0274 | 0.0183 | 0.1472 | 0.0623 |
| Cg1 | 0.4810 | 0.0122 | 0.7596 | 0.7007 | 0.9207 | 0.8064 | 0.4029 | 0.7327 | 0.0681 | 0.0917 | 0.2090 | 0.0679 |
| PrL | 0.4430 | 0.1463 | 0.7629 | 0.9715 | 0.3466 | 0.1055 | 0.4049 | 0.9451 | 0.1401 | 0.0202 | 0.2185 | 0.3259 |
| IL | 0.1346 | 0.2449 | 0.7533 | 0.1295 | 0.1330 | 0.0077 | 0.3939 | 0.1288 | 0.0598 | 0.0106 | 0.1898 | 0.2641 |

#### dHPC damage changes interactions among other regions

As the comparison above focused on the nodes, we next examined differences between the dHPC and SHAM-nH networks based on their edges (correlation coefficients). First, we compared the distribution of correlations of each matrix between groups using a two-sample KS test. We observed significantly different correlation coefficient distributions between dHPC and SHAM-nH networks in all thresholds (**threshold 0.05:** D_151_ = 0.2527, p = 0.0125; **0.025:** D_106_ = 0.3396, p = 0.0042; **0.01:** D_67_ = 4795, p = 0.0005; **Figure 7a-c**). Next, we compared each correlation coefficient between the groups. We computed the Z-score of the group difference for each correlation coefficient and considered a score of |2| to be significant within the distribution. We observed 21 correlation differences with Z-scores above |2| (**Figure 7b**). In nearly 2/3 of the significant differences (15 out of 21), the stronger correlation coefficients belonged to the SHAM-nH network, and 9 of them belonged to SHAM-NH hubs in that threshold; whereas only 6 differences the stronger correlation coefficient belonged to the dHPC network, one of which belonged to a hub (**Figure 7c**). These results were similar across thresholds. In the 0.025 threshold, 19 out of 26 differences were higher in the SHAM-nH network (3 belonging to SHAM-nH hubs; **Figure 7b**), and in the 0.01 threshold, 20 out of 28 differences were higher in SHAM-nH network (9 belonging to SHAM-nH hubs; **Figure 7c**). Overall, these results show that the SHAM-nH network presented a higher number of significantly stronger correlations compared to the dHPC network, many of which belonged to SHAM-nH hubs for that threshold.

**Figure 7:**
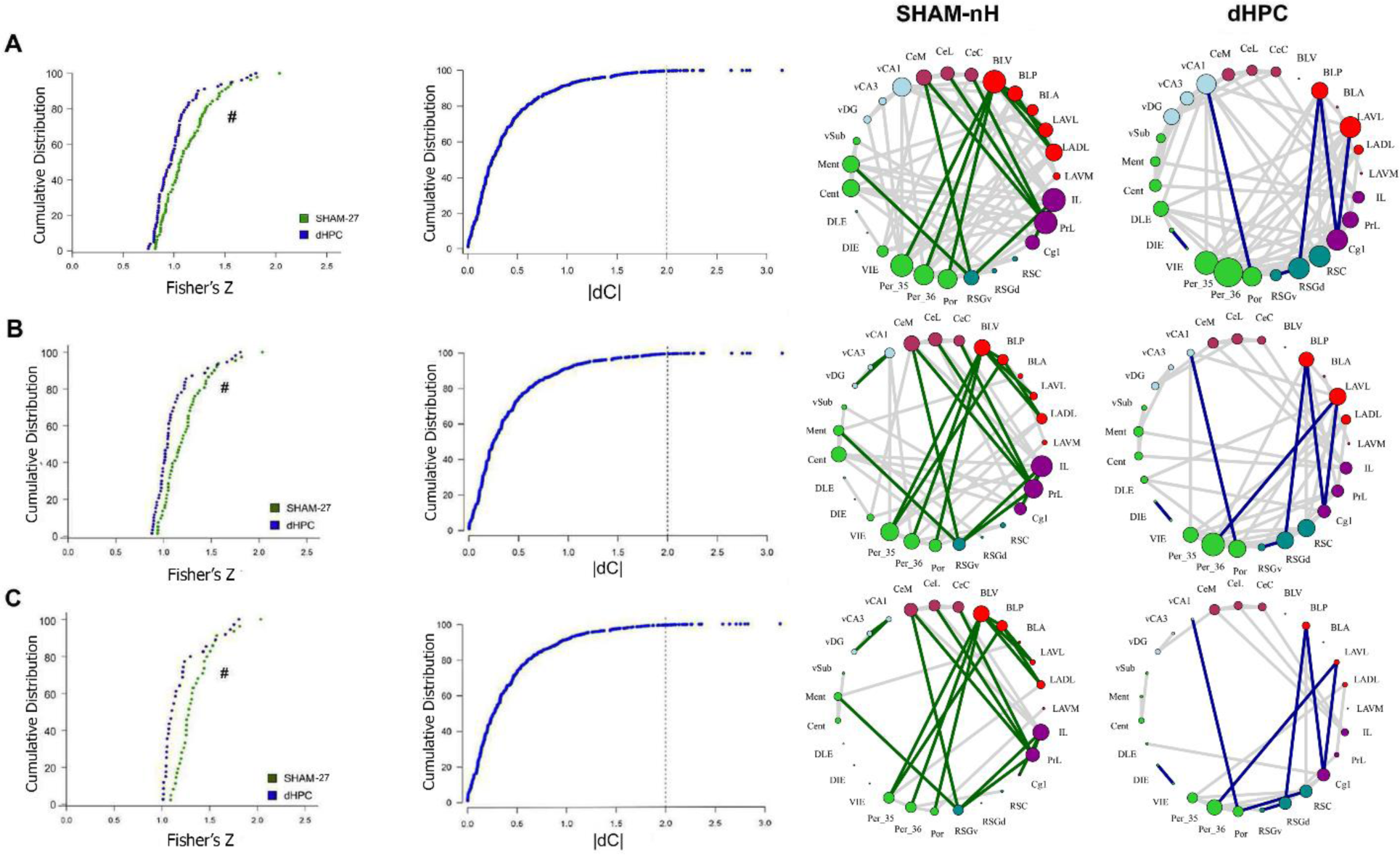
Connectivity Change in dHPC network. (A) Cumulative distributions of the Fisher’s Z transformed correlation coefficients from the SHAM-nH and dHPC matrices. The “#” indicates that these distributions are significantly different (Kolgomorov-Smirnov test, p<0.05). (B) Cumulative distribution showing the z-score of the correlation coefficient differences between the groups in each cell. The dashed line shows the absolute Z score of 2, revealing the values considered significant (beyond it) at the level of α = 0.05. Next, (C) the significantly different coefficients were plotted in each network, showing the network and nodes to which it belonged. The same procedure was performed in the 0.025 (D) and 0.01 (E) threshold networks, with similar results.

The different correlation distributions and the differences in correlation strengths between the networks add support to the hypothesis of different connectivity patterns in the dHPC network. Further, it suggests that dHPC indirectly influences interactions between other regions, most of which were observed to be weakened.

### Damaging the dHPC network Hubs

The network analysis revealed some differences between the dHPC and the SHAM (or SHAM-nH) networks. Particularly, the alternative hubs emerging in the dHPC network (Per_36 and RSC) and their statistically higher centralities compared to the SHAM-nH network suggest that these regions may increase in their importance to CFC learning in the absence of hippocampus. We empirically tested this hypothesis in the next two experiments by damaging both the dHPC and one of these hubs pre-training to CFC. Our hypothesis is whether further insult to the network would compromise the necessary structure of the network to promote CFC learning.

### Experiment 2 – Pre-training dHPC-Per double lesion does not impair CFC

In *Experiment 2*, because it was technically difficult to damage specifically the Per_36 and most animals had a significant part of the Per_35 damaged, we considered animals with lesions extending to both Per_36 and Per_35, denominating it Per. Henceforth, Per will be mentioned when Per_35 and Per_36 are considered together. During histological analysis, we excluded four rats from the dHPC-Per, two from the Per and one from the dHPC groups due to either extensive bilateral lesions to the regions surrounding Per (Temporal, Auditory, Parietal, Visual cortices, ventral CA1 or Lateral Amygdala), or no detectable dHPC and/or Per cellular loss in most slices examined. The final sample in this experiment was 38 (SHAM, dHPC and Per: N = 10/each; dHPC-Per: N = 8). In the remaining sample, cellular loss was mostly confined to the Per_36, Per_35 and to dHPC. In the dHPC and dHPC-Per groups, slight occasional damage was observed in the secondary Visual and Medial Parietal cortices overlying dHPC due to needle insertion (**Figure 8a**). In the behavioral analysis, the bootstrapped ANOVA showed no group difference (F = 0.842, p = 0.479; **Figure 8b-c**). The KS test showed no significant differences among groups’ distributions and the Cohen’s d values did not show any considerable effect size (**Figure 8** **bottom**). These results indicate that neither Per or dHPC-Per lesions affect CFC learning and memory.

**Figure 8:**
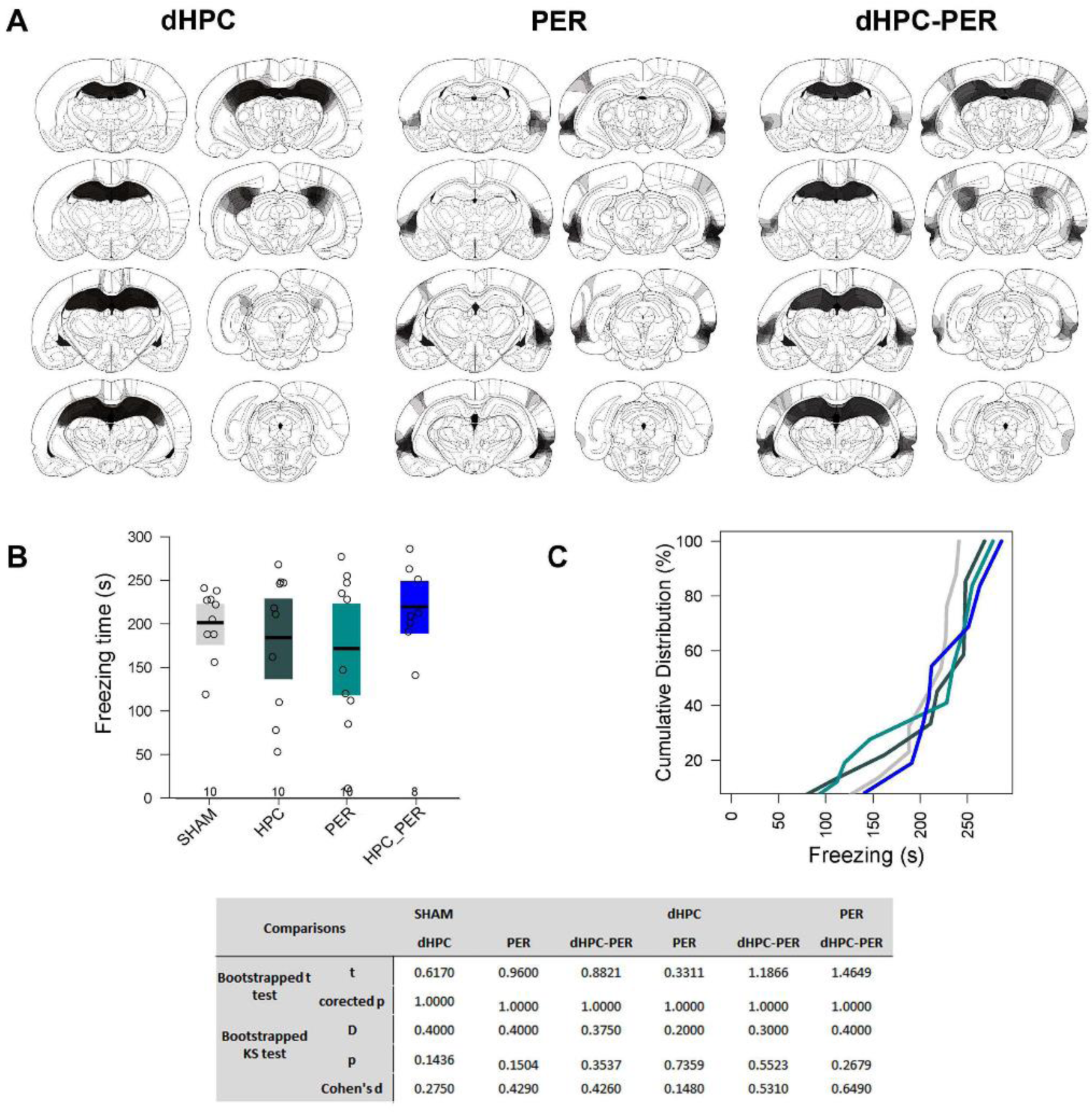
Per and dHPC-Per lesions on CFC learning. (A) Histological diagrams showing the distribution of areas damaged in dHPC, Per and dHPC-Per groups. The more overlapped the damaged areas across subjects, the darker the area. Mean and bootstrapped 95% IC of the total freezing time in SHAM, dHPC, Per and dHPC-Per groups during 5 min of CFC memory test. Dots show the sample distribution of each group. (C) Cumulative distribution of the total freezing time in each group in the same CFC memory test. The bottom table shows all the statistical tests performed and the corrected p-value for each comparison.

Previous studies observed no pre-training Per lesion effect on CFC (Phillips and LeDoux, 1995; Herzog and Otto, 1998), despite some contradictory evidence (Bucci et al., 2000). Our results support the hypothesis that pre-training Per and dHPC-Per lesions do not affect CFC learning and memory.

### Experiment 3 – Pre-training dHPC-RSC double lesion impairs CFC

During histological analysis, three rats from the RSC and one from the dHPC-RSC group were excluded from the analysis due to non-detectable cellular loss in most slices. The final sample in this experiment was 39 (SHAM: N = 10, dHPC and RSC: N = 9/each, dHPC-RSC: N = 11). The lesions affected mainly the dHPC and RSC, with frequent lesions to RSGd and occasional minor unilateral lesions of RSGv and secondary visual cortex. In the behavior analysis, the bootstrapped ANOVA revealed a main effect of group (F_3,35_ = 3.691, p = 0.01975), which the p-corrected t tests showed to be due to a lower freezing in the dHPC-RSC compared to that of the SHAM group (t_20_ = 3.315, p = 0.0270; **Figure 9**). No other significant differences were observed. This result was further confirmed by the KS test, which revealed significantly different distributions between the dHPC-RSC and the SHAM samples (D = 0.609, p = 0.0303). No other differences were observed (SHAM vs dHPC: D = 0.378, p = 0.330; SHAM vs RSC: D = 0.367, p = 0.377; dHPC vs RSC: D = 0.333, p = 0.316; dHPC vs dHPC-Per: D = 0.485, p = 0.098; Per vs dHPC-Per: D = 0.374, p = 0.289). The Cohen’s d values also confirmed the above results showing a large effect size between SHAM and dHPC-RSC means (d = 1.469). Lesser effect size values were observed in the other comparisons (SHAM vs dHPC: d = 0.463; SHAM vs RSC: d = 0.75; dHPC vs Per: d = 0.338; dHPC vs dHPC-Per: d = 1.056; Per vs dHPC-Per: d = 0.598), although the effect size between dHPC and dHPC-RSC was somewhat large.

**Figure 9:**
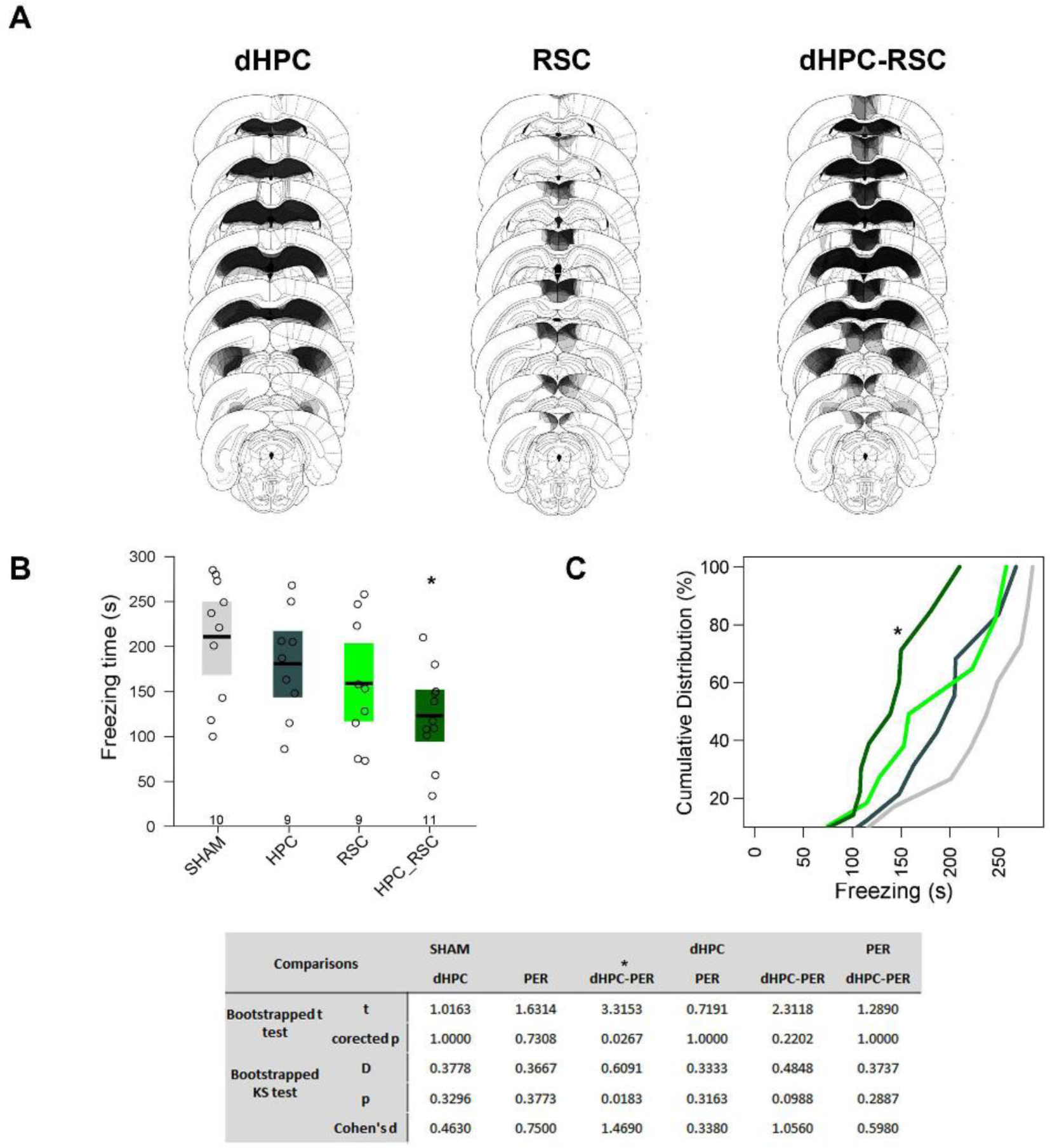
RSC and dHPC-RSC lesions on CFC learning. (A) Histological diagrams showing the distribution of areas damaged in dHPC, RSC and dHPC-RSC groups. The more overlapped the damaged areas across subjects, the darker the area. Mean and bootstrapped 95% IC of the total freezing time in SHAM, dHPC, RSC and dHPC-RSC groups during 5 min of CFC memory test. Dots show the sample distribution of each group. (C) Cumulative distribution of the total freezing time in each group in the same CFC memory test. The bottom table shows all the statistical tests performed and the corrected p-value for each comparison. “*” shows significant differences relative to SHAM group (corrected-p <0.05).

These results show that both dHPC and RSC contribute to CFC learning, although single lesion of these regions was not sufficient to impair CFC. Further, it supports the network analysis in *Experiment 1* that RSC becomes a key region in the dHPC network engaged in CFC learning.

## DISCUSSION

The present study employed network science to investigate CFC learning in dHPC-damaged rats. A fair amount of studies have observed CFC learning in absence of dHPC (Wiltgen et al., 2006; Fanselow, 2010; Zelikowsky et al., 2012), but no evidence had been provided, so far, regarding how a compensation mechanism might occur. Our study shows four main findings. First, we found that the CFC learning network under dHPC damage did not affect the small-worldness observed in the SHAM and SHAM-nH networks, and presented comparable levels of global and local efficiencies to the SHAM network. Second, we identified different hubs in each network, which were associated with different centrality levels between the dHPC and SHAM-nH networks. Third, differences in correlation coefficients distribution and strength suggest that dHPC indirectly influence interactions in the network. Fourth, by damaging the regions identified as hubs in the dHPC network, we showed that double lesion of dHPC and RSC, but not dHPC and Per, disrupt CFC learning and memory. Overall, despite the unaltered topology, dHPC network was sufficiently different such that alternative hubs emerged.

Many studies have observed small-world architecture in both anatomical and functional brain networks (Bassett and Bullmore, 2006; 2016). Small-world architecture is proposed to confer optimized cost-efficiency to interactivity (Achard and Bullmore, 2007) and protection to central regions to targeted attack, when compared to other topologies (i.e. scale free networks; Achard et al., 2006). Further, topology (integration and segregation) have also been observed to coordinate network reconfigurations during learning (Bassett et al., 2011), recollection (Fornito et al., 2012) and to predict errors in learning tasks (Ekman et al., 2012), providing evidence of the importance of topology to cognitive function. In agreement with this framework, the present data show unaltered CFC memory behavior and network integration and segregation levels, maintaining small-worldness in CFC learning network even under dHPC insult.

In the present study, the RSC and Per_36 showed stable hubness in the dHPC network and presented higher centrality levels compared to SHAM-nH network. These regions are deemed as central components of the proposed antero-temporal (AT, Per_36) and postero-medial (PM, RSC) memory systems that converge to the hippocampus (Ritchey et al., 2015), suggesting that dHPC damage increases the importance of the ‘upstream’ regions. Albeit the validation experiments showed impaired CFC memory only in the dHPC-RSC double lesions, but not dHPC-Per, the centrality comparison supports this double lesion data. The RSC displayed more robust centrality differences, with significance in more metrics and in all thresholds. These stable centrality differences may be reflecting an increased demand over – and dependence on – the RSC in the dHPC network.

Our data also corroborates the current framework of Per and retrosplenial cortex (RSG) functions. The Per is related to recognition, affective processing and associative memory of non-spatially referenced cues (Kealy and Commins, 2011; Suzuki and Naya, 2014; Kinnavane et al., 2017), whereas the RSG is important for processing spatial, contextual information and episodic memory (Ritchey et al., 2015; Todd and Bucci, 2015). Therefore, it is parsimonious that the CFC network under dHPC damage be more dependent on RSG than on Per.

The RSG has been considered an anatomical connector of the diencephalon, medial temporal lobe and cortices implicated in anterograde amnesia (Aggleton, 2008; Vann et al., 2009). A recent re-emerged interest in the RSG provided a diverse number of evidences highlighting its function. For instance, studies in humans showed increased activity in RSG for stable landmarks when navigating in virtual reality environments (Auger et al., 2012; Auger et al., 2015, 2017). Studies in animal models provided evidence that RSG integrates, encodes and stores spatial information (Czajkowski et al., 2014; Alexander and Nitz, 2015; Jacob et al., 2017), that it is necessary during spatial navigation (Pothuizen et al., 2008; Nelson et al., 2015a) and context fear learning and memory (Keene and Bucci, 2008b; Cowansage et al., 2014; Todd et al., 2017). This framework suggests that the RSG is an important component of spatial learning and memory systems. Furthermore, RSG is highly interactive with regions known to be involved in spatial and contextual learning such as hippocampus (Cooper and Mizumori, 2001) and Por (Robinson et al., 2012). The present results are in line with these findings and suggest that in dHPC absence, contextual learning networks might increase their dependence over the RSG.

On a different perspective, increased functional connectivity and centrality levels are hallmarks in patients with traumatic brain injury (TBI; Hillary et al., 2015). Some authors propose that hyperconnectivity is a natural response to brain insult and reflects an overload of alternative reminiscent pathways still capable of supplying the cognitive demand (Caeyenberghs et al., 2016; Hillary and Grafman, 2017). Regions exhibiting increased connectivity generally compose network rich clubs, including the RSG and Per (reigons defined as PCC, and ParaHipp, respectively; Hillary et al., 2014; Hillary et al., 2015). The present findings extend the occurrence of the increased centrality levels of RSG and Per after CFC learning under dHPC damage and provide evidence for the validity of the network measures, under some stability of the effect.

The hyperconnectivity accounts also challenges the notion that post-lesion connectivity increases are an adaptive compensatory mechanism. For instance, increased connectivity was observed to not be predictive of cognitive performance and even to diminish in the pre-frontal cortex after sustained practice of a working memory task (Medaglia et al., 2012). Whilst much is unknown about post-injury increased functional connectivity, future work could test if remote CFC memory or multiple CFC sessions result in a dHPC network closer to that of controls.

We also observed an indirect influence of dHPC lesion on interactions among other regions, which is consistent with both simulation of functional brain activity under brain damage (Alstott et al., 2009), and studies on unilateral focal brain lesions (Corbetta et al., 2005; He et al., 2007). This non-local alteration in connectivity was associated with behavioral impairments in patients. Although we did not observe a contextual fear memory impairment, the altered pattern of connectivity observed gives support to a partially different CFC learning network under dHPC damage, and suggests that what is learned (associated to the shock) might be different under lesion.

Importantly, the lack of effect on pre-training lesions involving Per should not be taken as evidence against its involvement in CFC. As RSC and Per in this study, pre-training hippocampal lesions do not impair CFC either (Wiltgen et al., 2006). Further, post-training lesions to all these regions resulted in impaired CFC memory (Bucci et al., 2000; Burwell et al., 2004; Wiltgen et al., 2006; Todd et al., 2017) and after pharmacological manipulations (Schenberg and Oliveira, 2008; Albrechet-Souza et al., 2011; Corcoran et al., 2011), evidencing that these regions do play a role in CFC. Our hypothesis was focused on whether compensation would still occur further targeted network damage to the dHPC network.

Moreover, previous studies employing pre-training single lesions on both Per and RSG have reported conflicting results regarding their effect on CFC. On Per lesions, one study reported impaired CFC memory in Per-damaged animals (Bucci et al., 2000), whereas other reports did not find impairment (Phillips and LeDoux, 1995; Herzog and Otto, 1998). These studies employed different lesion methods and behavioral parameters, rendering it difficult to point a source of the discrepancy. Although the present study employed methods closer to that of Bucci and colleagues (2000), the conflicting results remain. Regarding RSG, Keene and Bucci (2008b, a) have consistently observed impaired CFC memory in pre-training RSG lesions, whereas another study did not find such impairment (Lukoyanov and Lukoyanova, 2006). Our procedures were as similar as possible to that of Keene and Bucci (2008b), however, we aimed for the RSC instead of the whole RSG. Although we did damage portions of RSGd in some animals it is possible that our lack of effect on RSC single lesions was due to not damaging the whole RSG. Alternatively, it is possible that Per and RSG single lesions may be at least partially compensated just as dHPC lesions, resulting in higher rates of mixed results due to a less effective learning (Fanselow, 2010). Despite the unimpaired behavior in dHPC-damaged animals, it is very likely that the contextual information learned is different (Frankland et al., 1998; Nadel, 2008). Some authors discussed about the complexity of the CS under hippocampal damage (Rudy, 2009; Fanselow, 2010), however, clearly assessing the content learned as CS in CFC preparations remains as a limitation. Findings from tasks that allow a better assessment of the learned content strongly suggest that both Per and RSG support configural learning – defined as complex stimuli bound together in a stimulus-stimulus manner. For instance, Per-damaged rodents have impaired complex visual discrimination tasks (Eacott et al., 2001; Hales et al., 2015), and RSG-damaged rodents have impaired spatial memory in tasks in which spatial cues moved between trials (Hindley et al., 2014; Nelson et al., 2015b). Further, RSG was shown to integrate distributed spatial information across delimiting marks (Alexander and Nitz, 2015). These data suggest that RSC and Per can support some configural learning in dHPC-damaged animals. This is supported by studies employing whole-hippocampus damage and complex maze tasks (Winocur et al., 2010).

### Methodological considerations and Limitations

There are some points about the present study that need attention when interpreting the results. First, the lesion method used in *Experiment 1* (electrolytic lesion) does not spare fibers of passage, which may have affected connections between other regions. Whilst this could have altered the network more than intended, the behavior data suggests that the network is likely to contain the elements required in CFC learning and memory since no impairment was observed. Furthermore, the networks studied here, which are based on pCREB expression, identified similar hubs to recent anatomical studies based on larger tract-tracing databases (Binicewicz et al., 2015; Bota et al., 2015), making a confounding effect of fiber lesion unlikely.

Second, the *Experiment 1* differs from *Experiments 2* and *3* in number of shocks during the training session. Single shock CFC sessions is generally a weaker experience and tend to yield more variable levels of behavior. We used the three shocks procedure to ensure a robust performance level in *Experiments 2-3* such that impairments would be more detectable. Additionally, the performances of SHAM controls and dHPC groups were very similar, ruling out the possibility of a ‘hidden’ memory impairment in the dHPC group in *Experiment 1*.

### Conclusion

There is growing interest in the use of network approaches to predict cognitive performance from brain imaging data (Bassett et al., 2011; Ekman et al., 2012; Fornito et al., 2012). However, formally testing predictions in human experimentation is still a challenge called for attention (Petersen and Sporns, 2015). We applied network analysis in rodent models such that we could empirically test the validity of these models later. We found that new hubs identified in the CFC network under dHPC damage may compromise the formation of the functional network necessary for CFC learning and memory. Future employment of finer techniques (i.e. optogenetics, transgenic animals) may provide sophisticated ways to test network predictions.

## MATERIALS AND METHODS

### Subjects

A hundred and thirty nine male Wistar rats weighting 300-370g were obtained from the university vivarium (CEDEME, SP). They were housed in groups of 4 – 5 and maintained on a 12h light/dark cycle, room temperature of 22 ± 2°C, with free access to food and water. All experiments were approved by the University Committee of Ethics in Animal Research (#0392/10, #409649 and #7683270116) and were in accordance with National Institutes of Health Guide for the Care and Use of Laboratory Animals.

### Surgery

The rats were anesthetized with Ketamine (90mg/kg, Ceva, Paulínia, Brazil) and Xilazine (50mg/kg, Ceva, Paulínia, Brazil), and mounted into a stereotaxic frame (David Kopf Instruments, Tujunga, CA). Each animal had their scalp incised, retracted and the bregma and lambda horizontally adjusted to the same plane. Small holes were drilled in the skull in the appropriate sites. The rats received bilateral electrolytic lesions in the dHPC by an anodic current (2 mA, 20 s) passed through a stainless steel electrode insulated except for about 0.7 mm at the tip. The following coordinates were used: −4.0 mm from bregma (AP), ± 2.0 and ± 4.0 mm from the midline (ML) and −3.6 mm from the skull surface (DV). Control (SHAM) animals underwent the same procedure except that they did not receive currents. After the surgery, the rats received antibiotic and diclofenac intramuscularly (3mg/kg, Zoetis, Madison, NJ) and were allowed to recover for 15 days. To avoid corneal lesions associated to the anesthetic used, the rats had their eyes hydrated with ophthalmic gel (Bausch & Lomb, Rochester, NY) and received a post-surgery injection of yohimbine (2mg/kg, Sigma, St. Louis, MO).

In *Experiment 2* the surgeries were performed as above, but the rats received bilateral neurotoxic lesions in the dHPC, Perirhinal cortex (Per), both (dHPC-Per) or SHAMs. The lesions were made by N-methyl-D-aspartic acid (NMDA, 20 mg/ml in 0.1 M phosphate buffered saline, pH 7.4; Sigma, St. Louis, MO) injected by a 10 µl Hamilton syringe held by a microinjector (Insight, Ribeirão Preto, Brazil) and connected to 27 gauge injecting needles by polyethylene tubes. In the dHPC, 0.45 µl of NMDA was injected at a rate of 15 µl/min in each of the following coordinates: (1) AP: −2.8 mm, ML: ± 1.5 mm and DV: −3.6 mm; (2) AP: −4.2 mm, ML: ± 1.5 and ± 4.0 mm and DV: −4.0 mm. In the Per, 0.1 µl of NMDA was injected (0.1 µl/min) in each of the following coordinates: AP: −2.6, −3.5, −4.4, −5.4 and −6.5 mm, ML: ± 5.9, ± 6.1, ± 6.1, ± 6.5 and ± 6.4 mm, DV: −7.4, −7.4, −7.4, −7.2 and −7.0. The needle remained in place for an additional 3 min. The post-surgical procedures were identical to those in Experiment 1.

In *Experiment 3*, surgeries were performed as in *Experiment 2*, but for lesions of the dHPC, disgranular retrosplenial (RSC), both (dHPC-RSC) or SHAMs. In the RSC, 0.2 µl of 20 mg/ml NMDA was injected (0.1 µl/min) in the following coordinates: AP: −3.0, −4.0, −5.0, −6.0 and −7.3 mm, ML: ± 0.4, ± 0.4, ± 0.5, ± 0.7 and ± 0.8 mm, DV: −0.8, −1.0, −1.0, −1.1 and −1.5 mm.

### Apparatus

We used a fear conditioning chamber (32 x 25 x 25 cm, Med Associates, St. Albans, VT) equipped with Video Freeze System. The chamber was composed of aluminum (sidewalls), polycarbonate (front wall and ceiling), white opaque acrylic (back) pieces and a grid floor of stainless steel rods (4.8 mm thick) spaced 1.6 cm apart. A sound-attenuating chamber with fans (60 dB) provided background noise and white house lights enclosed the chamber. After each animal, the chamber was cleaned with 10% ethanol.

### Contextual Fear Conditioning (CFC)

Before every experiment, all animals were gently handled for 3 consecutive days.

In *Experiment 1*, during the training session, the rats were individually placed into the conditioning chamber for 2 min, received a 1 s, 0.8 mA footshock, and were returned to their homecage after 1 min. One additional control group of SHAM animals (Imm) was placed in the conditioning chamber, received an immediate footshock and was immediately returned to the homecage. Half of the cohort was re-exposed to the context 48h later for 5 min to test contextual fear memory. Behavior was recorded in both sessions by a micro-camera in the chamber. An experimenter blind to the grouping measured the freezing behavior, defined as complete immobility except for breathing movements (Bouton and Bolles, 1980), which served as our measure of contextual fear memory.

In *Experiments 2 and 3*, rats were placed into the conditioning chamber for 2 min, but received <u>three</u> 1 s, 0.8 mA footshocks, with 30 s inter-trial interval, and were returned to their homecage after 1 min. The rest of the procedure is identical to *Experiment 1*, except that there was no Imm control group.

### Perfusion and Immunohistochemistry

Phosphorylated CREB (pCREB) has a two-phase peak expression profile, which the latter (3-6 h) was shown to present a clearer associative learning-specific expression (Stanciu et al., 2001; Trifilieff et al., 2006). Therefore, we used a 3h time window of pCREB expression in our study. Three hours following training in *Experiment 1*, half the cohort was deeply anesthetized and perfused transcardially with buffered saline and 4% paraphormaldehyde (PFA) in 0.1 M sodium buffer (pH 7.4). The brains were extracted, post-fixed in PFA, cryoprotected in 20% buffered sucrose, frozen and stored at −80°C. The brains were coronally sectioned in 30 μm thick slices in a cryostat (®Leica, Wetzlar, Germany) and stored in 4 serial sets. One set was collected in glass slides and stained with cresyl violet for morphological and lesion analysis, another set was used for phospho-CREB immunolabelling and the two remaining were stored for future studies. Immunolabelling was performed in free-floating sections using anti-phospho-CREB (1:1000, Santa Cruz, Dallas, TX) as primary rabbit polyclonal antibody. A Biotinylated goat anti-rabbit antibody (1:800, Vector Labs, Burlingame, CA) was used as secondary antibody. The reaction was revealed using the avidin-biotin peroxidase method conjugated to diaminobenzidine as the chromogen (ABC and DAB kits, Vector Labs, Burlingame, CA) as described previously (de Oliveira Coelho et al., 2013).

### pCREB quantification

The pCREB expression was measured in 30 brain regions including hippocampal, parahippocampal, amygdalar and prefrontal regions (see Table **1**) previously shown to have involvement in FC and/or context learning. The dHPC group had 27 regions measured, since dCA1, dCA3 and dDG were damaged. The regions were delimited manually using ImageJ free software. The anatomical delimitation was based on the Rat Brain Atlas Paxinos and Watson (2007) as on other anatomical studies (see Table **1**; Insausti et al., 1997; Burwell, 2001; Sugar et al., 2011). Images (32-bit RGB) were taken at 4X and 10X magnifications using a light microscope (Olympus, Waltham, MA), and pCREB-positive cells quantified using the automated, high-throughput, open-source CellProfiler software (Carpenter et al., 2006). A pipeline was created to calculate the area of each region in mm^2^ and to identify stained nuclei based on their intensity, shape and size (20-150 μm^2^; **Figure 2**). The quantification was performed bilaterally in 6 sections/region (3 in each hemisphere). The data was expressed in nuclei density (nuclei/mm^2^). In each region and animal, three sections quantified bilaterally were averaged and computed as the expression data.

The pCREB is known to possess both a higher baseline and a higher expression profile (around twofold) compared to c-fos, an immediate early more commonly used as a proxy for neuronal activity (Hall et al., 2001; Stanciu et al., 2001; Colombo et al., 2003). Although a baseline signal close to zero is preferable in most studies, for correlation-based connectivity inference it blunts sensitivity to observe negative correlations, as a diminished expression is less observable. Detecting possible negative correlations was desired in our study, making pCREB a suitable proxy for neuronal activity. Further, pCREB has a well distinguishable expression in associative learning studies (Stanciu et al., 2001; Colombo et al., 2003; Trifilieff et al., 2006).

### Histology

In all experiments, the histological examination of the lesions was performed in the cresyl violet stained slices (150 µm apart) using a light microscope (Olympus, Waltham, MA). Lesions were identified visually as presence of tissue necrosis, absence of tissue or marked tissue thinning. Animals with no bilateral lesions of the target region or with lesions present in less than half the slices analyzed were excluded. An expressive bilateral lesion (50%) of untargeted regions was also an exclusionary criterion.

### Functional Connectivity and Network Generation

Different from the typical neuroimaging studies in humans, which acquire multiple measurements across time (i.e. EEG, fMRI), task-dependent large-scale brain activity in experimental animals is more limited. As immunohistochemistry provides a single *post-mortem* measure per region per animal, inter-regional co-activation is assessed across subjects. We used the pCREB-positive nuclei density to compute the Pearson correlation coefficient between all possible pairs of regions in each group (total of 435 coefficients in SHAM and 351 in the dHPC group). As SHAM matrix has 3 regions (dCA1, dCA3 and dDG) more than dHPC matrix, a “SHAM with no dorsal hippocampal regions” (SHAM-nH) was also calculated. The network derived from this matrix served to directly compare the network of these groups. Three thresholds were applied to the correlation matrices, maintaining only coefficients with two-tailed significance level of p ≤ 0.05, 0.025 or 0.01. This resulted in weighted undirected network graphs composed by the brain regions (nodes) and the remaining inter-regional correlations (edges), representing connections between the regions (**Figure 3**). The network analyses were performed in the networks of all thresholds.

### Network Measures

#### Topological metrics

This analysis was performed in distance (1 – Pearson’s r) matrices derived from the thresholded correlation matrices. We assessed the networks topology using global efficiency (Geff) as our measure of integration and mean local efficiency (Leff) as our measure of Segregation (Latora and Marchiori, 2001). Geff is defined as the mean of the inverse of all shortest paths in the network. And Leff is defined as the Geff applied to a subgraph composed by all neighbors of a given node.

Brain networks have been consistently characterized as possessing a *small-world* topology (Sporns and Zwi, 2004; Achard et al., 2006; Bassett and Bullmore, 2016). Small-worldness is usually estimated by metrics of integration and segregation, evincing equivalent integration and higher segregation relative to random networks (Watts and Strogatz, 1998). We compared the Geff and mean Leff of our empirical networks to those of randomized networks to test whether the empirical networks were small-world and if dHPC lesion affects the network small-worldness.

#### Centrality Metrics and Hub Identification

Hubs were identified using four centrality metrics: weighted degree (Wdg), eigenvector (Evc), Closeness (Clo) and Betweenness (Bet). For each metric, we intersected the 25% most central regions (upper quartile) of all four metrics and considered any regions within the intersection of at least three metrics as a hub for that threshold. To ensure a hub identification that was irrespective of thresholding, we intersected the hubs in each threshold and considered a stable hub any region present in all thresholds. As Wdg and Evc are connection-based metrics (based on number of connections), and Clo and Bet are distance-based metrics (based on short paths), we ensured that these regions were highly ranked in at least one metric of each type.

### Statistical Analysis

In the cohort tested for fear memory, we compared the Total Freezing Time during memory test between the groups by three statistical tests: one-way ANOVA, Kolgomorov-Smirnov (KS) tests and Cohen’s d effect size. In the ANOVAs and KS tests, we used a bootstrap resampling. The bootstrap resampling was defined by 1) randomly resampling the sample, with replacement of subjects by others (from the sample), 2) calculating the statistics of interest (i.e. F_resampled_) and 3) repeating it many times (10000). It generates an empirical sample-based artificial distribution of the statistics of interest under the null hypothesis, and allows to test if the empirical data statistics (F_empirical_) differs from random null hypothesis distribution. The p-value was calculated as the frequency of F_empirical_ occurring in the resampled distribution [p = (F_resampled_ > F_empirical_)/10000]. There is no normality (or any other) assumption to bootstrap resampling tests, allowing comparisons when the population distribution is not normal or unknown. Multiple comparisons were assessed by t tests with bootstrap resampling, as above, correcting the p-value by the number of concomitant comparisons.

In the cohort that had their brains immunolabelled for pCREB, the pCREB expression was quantified in 30 regions (27 in the dHPC group) as positive nuclei/mm^2^, and each region was compared between the groups using t tests with bootstrap resampling (as above), correcting the p-value with a false discovery rate (fdr) test (Benjamini and Hochberg, 1995).

In the hypothesis test for small-world network, each empirical network was ‘rewired’ as described previously (Maslov and Sneppen, 2002) to generate 10000 random, null hypothesis networks with the same number of nodes, edges, weights and degree distribution. Each network was rewired a number of times equal to half the number of their edges to generate the randomized networks. We calculated the Geff and mean Leff empirical/random ratio for each randomized network. It was expected for the Geff ratios to be around 1 and the mean Leff ratios to be above 1.

After the hub identification, we directly compared the centrality level of each region (in each threshold) between the dHPC and SHAM-nH networks using a permutation test. In the permutation procedure, we 1) randomized the grouping labels <u>without replacement</u>, 2) calculated the centrality values differences [Diff = C_SHAM_ − C_dHPC_] in each region and 3) repeated it 10000 times. The p-value was calculated as the frequency of the empirical difference (Diff_empirical_) occurring in the resampled (Diff_resampled_) distribution [p = (Diff_resampled_ > Diff_empirical_)/10000]. No comparisons with the SHAM network were performed as both networks have to be the same size. To test whether dHPC lesion influences interactions between other regions in the network, we compared the correlation coefficients between SHAM-nH and dHPC networks. We normalized the thresholded matrices using a Fisher’s Z transformation and compared the normalized correlation coefficient distributions in the dHPC and SHAM-nH networks with a two-sample KS test. Next, we calculated the z-score of the correlation coefficient difference between each cell of the matrices as in the formula bellow, defining an index of connectivity change, as done previously (Alstott et al., 2009). The Z-score values above |2| were considered significant (corresponding to a level of significance of α = 0.05). We verified which group possessed each significantly higher coefficient, and which nodes they connect.

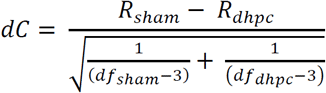

where df is the degree of freedom in each group.

In all analyses, a corrected-p < 0.05 was considered significant. All statistical and graph theory analyses and figures were performed in R studio (R, 2013) using custom-written routines (available at https://github.com/coelhocao/Brain_Network_analysis) and the packages igraph (Csardi and Nepusz, 2006), Matrix (Bates and Maechler, 2015), lattice (Sakar, 2008), ggplot2 (Wickham, 2009), corrplot (Wei, 2013), car (Fox and Weisberg, 2011) and VennDiagram (Chen, 2015).

## Acknowledgements

We thank André Fujita, André Cravo and Altay Lino de Souza for valuable comments and insights during the development of the study.

